# Integration of a *FT* expression cassette into CRISPR/Cas9 construct enables fast generation and easy identification of transgene-free mutants in Arabidopsis

**DOI:** 10.1101/663070

**Authors:** Yuxin Cheng, Na Zhang, Saddam Hussain, Sajjad Ahmed, Wenting Yang, Shucai Wang

**Affiliations:** Key Laboratory of Molecular Epigenetics of MOE, Northeast Normal University, Changchun, China; College of Life Science, Linyi University, Linyi, China

**Keywords:** CRISPR/Cas9, genome editing, bolting time, FT, AITR1

## Abstract

The CRISPR/Cas9 genome editing technique has been widely used to generate transgene-free mutants in different plant species. Several different methods including fluorescence marker-assisted visual screen of transgene-free mutants and programmed self-elimination of CRISPR/Cas9 construct have been use to increase the efficiency of genome edited transgene-free mutants isolation, but the overall time length required to obtain transgene-free mutants has remained unchanged in these methods. We report here a method for fast generation and easy identification of transgene-free mutants in Arabidopsis. By generating and using a single *FT* expression cassette-containing CRISPR/Cas9 construct, we targeted two sites of the *AITR1* gene. We obtained many early bolting plants in T1 generation, and found that about two thirds of these plants have detectable mutations. We then analyzed T2 generations of two representative lines of genome edited early bolting T1 plants, and identified plants without early bolting phenotype, i.e., transgene-free plants, for both lines. Further more, homologues *aitr1* mutants were successful obtained for both lines from these transgene-free plants. Taken together, these results suggest that the method described here enables fast generation, and at the mean time, easy identification of transgene-free mutants in plants.

## Introduction

Shortly after the CRISPR (clustered regularly interspaced short palindromic repeats) RNA-guided Cas9 (CRISPR-associated protein 9) endonuclease being reported to be able to cleave double-stranded DNA, therefore generate mutations in eukaryotic cells [1,2], CRISPR/Cas9 mediated genome-editing has been successful used for gene editing to generate mutations in several different plants including the model plant Arabidopsis and crops such as rice, tobacco and wheat [3–5]. Since then, the CRISPR/Cas9 genome-editing techniques including the double-stranded DNA cleaving based editing and the nucleotide substitution based editing have been widely used to generate mutations in different plant species, in some cases, to improve agronomic traits such as yield, quality and biotic and abiotic stress tolerances [6–12]. Thanks to their high efficiency in genome editing and the used of engineered Cas9 variants with expanded target space, CRISPR/Cas9 genome-editing systems have brought a bright future for plant breeding [6,12–14].

The presence of Cas9 T-DNA in CRISPR/Cas9 genome-edited mutants may affect the phenotypic stability and heritability of the mutation, and transgene-free is likely required for commercial application of CRISPR/Cas9 genome-edited crops [7, 10]. Therefore, isolation of transgene-free mutants is one of the most important steps for generation mutants by using CRISPR/Cas9 genome-editing. However, isolation of transgene-free mutants by using the traditional genetic segregation and backcross based genotyping is time consuming and laborious [7,8,10]. To improve the efficiency in transgene-free mutant isolation from CRISPR/Cas9 genome edited plants, a few different methods have been established [6–8,10]. These methods include the fluorescence maker-assisted selection, which allows to isolation transgene-free mutants based on the observation of the absence of fluorescence in seeds produced by transgenic plants [7]; the active interference element mediated selection, which allows herbicide-dependent isolation of transgene-free plants [8]; and the programmed self-elimination system, which allows only transgene-free male gametophytes to produce seeds [13]. All these methods greatly reduced workload for transgene-free mutant isolation. However, the overall time length required for the whole process of mutant generation, from plant transformation, to mutatant identification, and then transgene-free mutant isolation remained largely unchanged.

Appreciate flowering time is critical for successful sexual reproduction in flowering plants [15]. In order to achieve sexual reproductive successful, flowering plants need to sense and respond to environmental stimuli appropriately, and then integrate the environmental information with endogenous signals to make transit from vegetative growth to flowering [16–18]. Accumulated evidence suggest that flowering time in Arabidopsis is controlled by several different regulators, including CO (CONSTANS), SOC1 (SUPRESSOR OF CONSTANS OVEREXPRESSION1), FLM (FLOWERING LOCUS M), FLC, FLK (FLOWERING LOOCUS K HOMOLOGY (KH) DOMAIN), VRN2 (VERNALIZATION 2), MAF2 (MADS AFFECTING FLOWERING 2) and FT (FLOWERING LOCUS T) [15,17,19–24]. Among them, FLC and CO are major regulators involved in vernalization and photoperiod, the two most important environmental stimuli that control the switch from vegetative growth to flowering, respectively, and they function immediately upstream of FT to regulate the switch [15–17].

FT is the key positive regulator of flowering in Arabidopsis, and at least some of the FT homologues in other plants species including medicago, rice, soybean and trees like poplar and pear also function as activator of flowering [25–28]. Due to its important role in flowering promotion, FT has been successfully used to reducing juvenile phase of many plants, therefore accelerated the process for plant breeding [15,27,29].

Considering that early flowering phenotype caused by overexpression of *FT* in plants is easy visible, and the resulted short life cycle will accelerate mutants generation, integration of a FT expression cassette into CRISPR/Cas9 may enable fast generation and easy identification of transgene-free mutants in plants. In this study, we introduced a *GmFT2a* expression cassette into the *pHEE* CRISPR/Cas9 vector, and inserted two *sgRNA* expression cassettes to target the *AITR1 (ABA induced transcription repressor1*) gene, which encodes a novel ABA signaling and abiotic stress tolerance regulating transcription factor in Arabidopsis [30]. We successfully obtained detectable mutations in *AITR1* in early bolting T1 Arabidopsis transgenic plants, and obtained homologues transgene-free *aitr1* mutants from T2 plants with normal bolting phenotypes.

## Results

### Generation of a *FT* expression cassette-containing CRISPR/Cas9 construct

Overexpression of *FT* in plants promoted flowering, therefore shorten the juvenile phase of the transgenic plants even in trees [27]. If used in CRISPR/Cas9 mediated gene editing, the easy visible early bolting phenotype caused by overexpression of *FT* may serve as an assistant selection marker for easy identification of transgene-free mutants, whereas shorten in juvenile phase resulted by early flowering may reduce the length of the overall time required for generating genome edited mutants, thereby providing a method for fast generation and easy identification of genome edited transgene-free mutants.

Considering that overexpression of *GmFT2a* in both Arabidopsis and soybean promoted flowering in transgenic plants [31,32], we decided to integrate a *GmFT2a* expression cassette into CRISPR/Cas9 vector for gene editing. The full-length coding sequence of *GmFT2a* was synthesized, and cloned into to the *pUC19* vector to generate the *35S:GmFT2a* construct. The whole *35S:GmFT2a-nos* expression cassette was then PCR amplified and cloned into the *pHEE* vector [33], at the *pme1* site to generate the *FT* expression cassette-containing CRISPR/Cas9 vector *pHEE-FT* (Figure 1a). In this vector, the *FT* expression cassette was inserted within the T-DNA fragment of the *pHEE-FT* vector as an independent expression cassette, and the expression of *GmFT2a* was controlled by the strong double *35S* promoter, which enables that *GmFT2a* can be expressed in a high level in transgenic plants generated by using the *pHEE-FT* vector, therefore may led to an easy visible early flowering phenotype in the transgenic plants.

**Figure 1.**
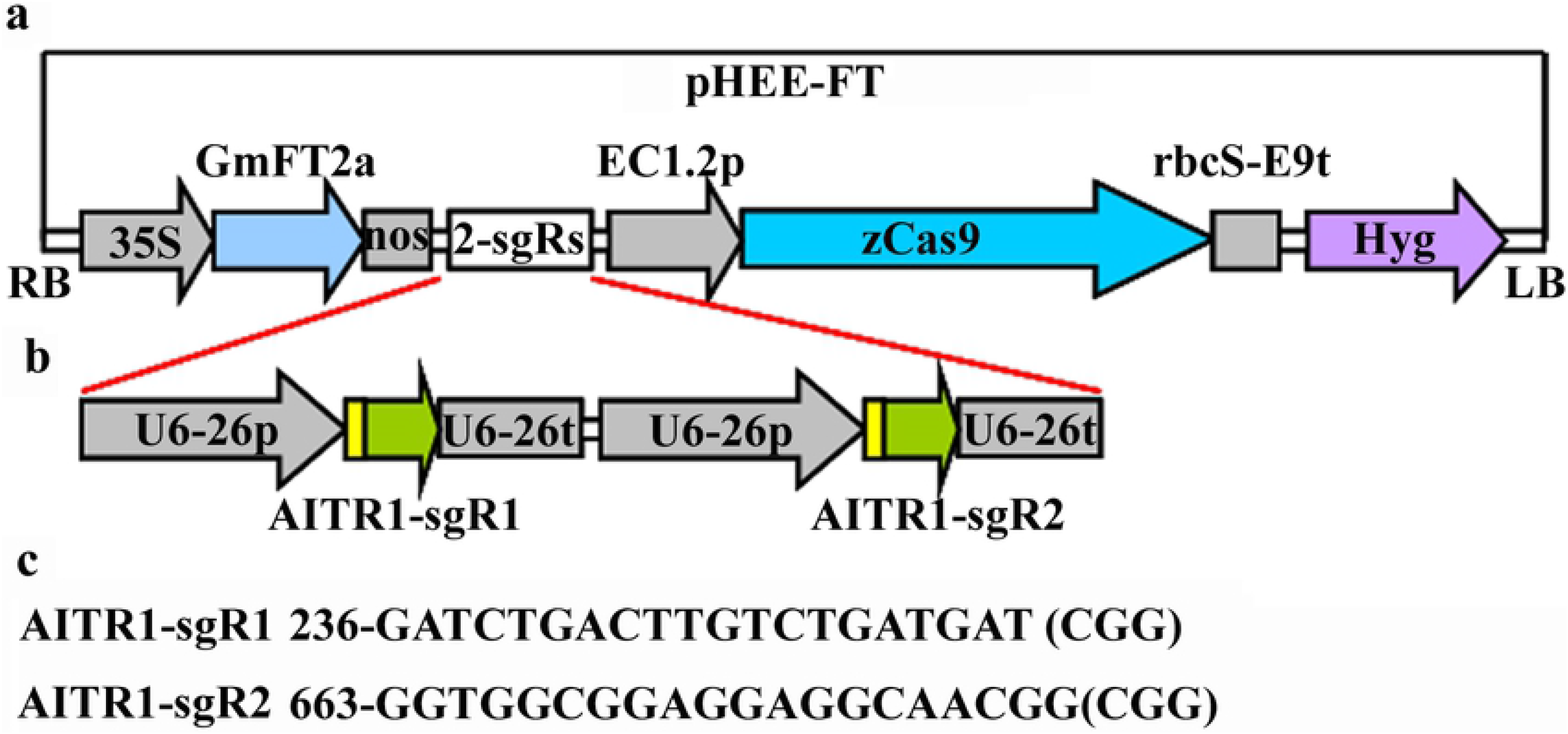
Generation of a *FT* expression cassette containing *pHEE* CRISPR/Cas9 construct for *AITR1* editing. (**a**) *pHEE* vector with a *FT* expression cassette. The full-length ORF sequence of *GmFT2a* was synthesized, and cloned into *pUC19* vector under the control of the *35S* promoter and terminated by the *nos* sequence. The *35S:GmFT2a-nos* cassette was amplified by PCR and then inserted into the *pHEE* vector at the Pme1 site by using Gibson assembly to generate *pHEE-FT* vector. (**b**) The *sgRNA* expression cassettes in the *pHEE-FT-AITR1* construct. The *sgRNA* sequences corresponding to the target sequences of *AITR1* were introduced into the *sgRNA* expression cassettes by PCR amplification, followed by Golden Gate reaction with the *pHEE-FT* vector. (**c**) Target sequences in *AITR1*. Numbers indicated the nucleotide position relative to the first nucleotide in the coding sequence of *AITR1*, PAM sites after the target sequences were indicated in the brackets.

### T1 transgenic plants generated using the *FT* expression cassette-containing CRISPR/Cas9 construct showed an early bolting phenotype

To examine if the genome editing using the *pHEE-FT* vector may enable fast generation and easy identification of transgene-free mutants, we made a *pHEE-FT* CRISPR/Cas9 genome editing construct to target *AITR1* (Figure 1b), a novel transcription factor gene that has been shown to regulate ABA signaling and abiotic tolerance in Arabidopsis [30], at two different target sites (Figure 1c).

After selected on antibiotic-containing plates, more than 70 T1 independent transgenic plants were obtained. As expected, early bolting phenotype was observed in the transgenic plants (Figure 2a). Bolting was observed in transgenic plants as early as 8 days after the seedlings were transferred from the antibiotic-containing plates into soil pots, and the bolting time for these T1 transgenic plants ranged from 8-19 days after the transfer, whereas that in the Col wild type plants ranged from 17-20 days after the transfer (Figure 2b).

**Figure 2.**
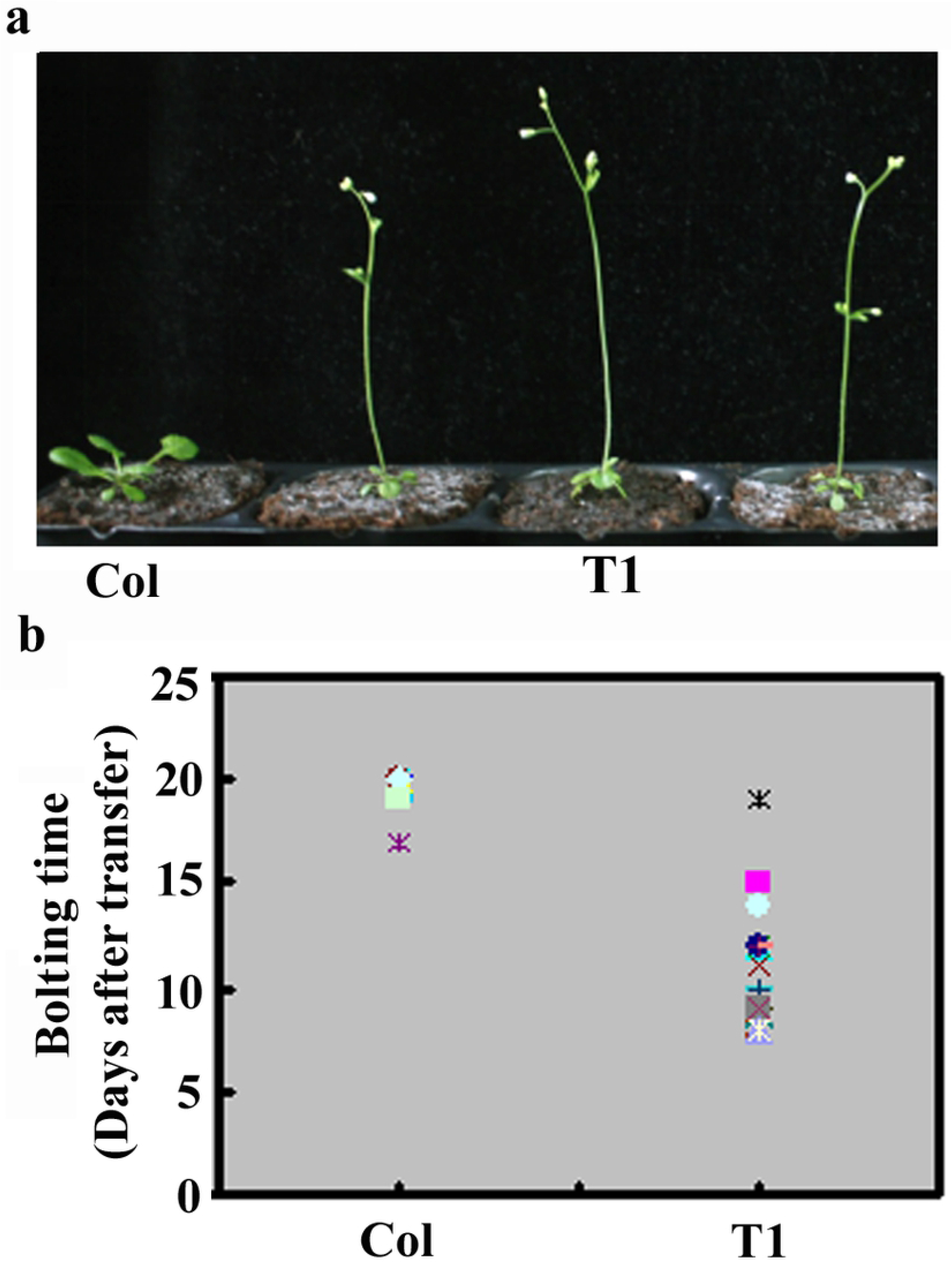
Early bolting phenotypes observed in the T1 transgenic plants. (**a**) Early bolting phenotype in some T1 transgenic plants. Transgenic plants were selected on antibiotic-containing 1/2 MS plates, and ~5-day-old transgenic seedlings were transferred into soil pots and grown in a growth room. As a control, seeds of Col wild type were germinated on 1/2 MS plates, and seedlings ~5-day-old were transferred into soil pots. Pictures were taken 10 days after the transfer. (**b**) Bolting time of the T1 transgenic plants. The date of bolting for the plants was recorded, and days after the transfer were calculated. For Col wild type plants, n=11. For transgenic plants, n=53.

### Homozygous genome edited mutants were obtained in the T1 transgenic plants

The *pHEE-FT-AITR1* CRISPR/Cas9 construct was made to target two sites in *AITR1*, therefore fragment deletion should be expected in the transgenic plants if both sites can be edited. Because we intended to use early bolting phenotype as an assistant marker for transgene-free mutant isolation, transgenic plants with an early bolting phenotype, i.e., bolted 8 or 9 days after transferred, were chosen for fragment deletion examination by PCR. Yet we included a few plants with medium or normal bolting time in this experiment to examine if this is a co-relationship between early bolting phenotype and genome editing status.

Indeed, smaller PCR product band was obtained (Figure 3a). However, among about 50 T1 plants examined, only two plants, i.e., lines 14 and 59 with early and medium bolting phenotype, respectively yield two PCR bands, but none of them produced only one smaller band (Figure 3b). All other plants produced only one larger PCR band with expected site for the full-length coding sequence of AITR1(Figure 3b). These results suggest that both target sites in *AITR1* can be edited, but homozygous fragment deletion mutants were not obtained in the plants examined.

**Figure 3.**
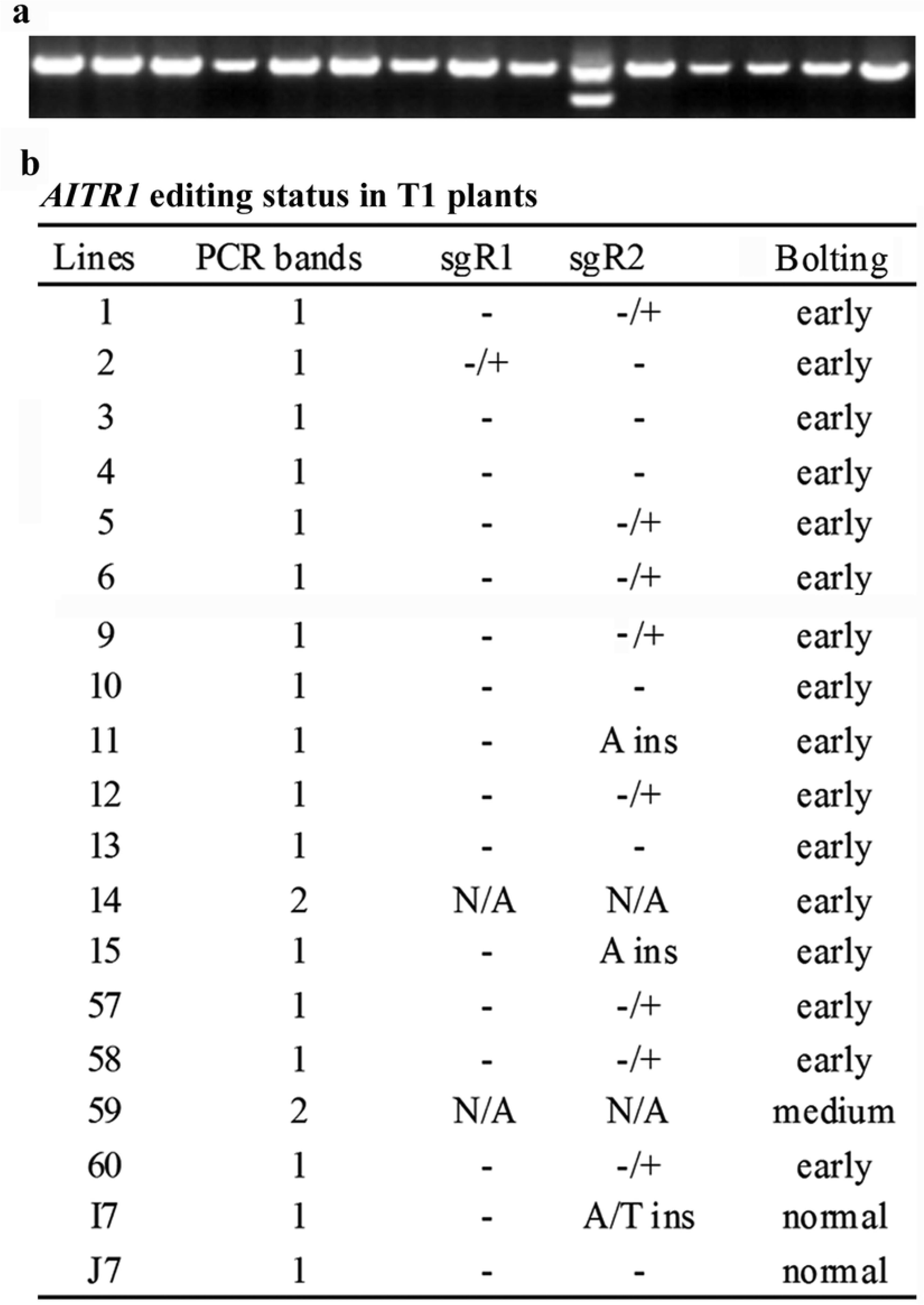
*AITR1* editing status in T1 transgenic plants. (**a**) PCR amplification of *AITR1* coding sequence in the T1 transgenic plants. DNA was isolated from leaves collected from individual T1 transgenic plants, and PCR was used to amplify coding sequence of *AITR1*. Picture is representative image of PCR results, showing the 1 or 2 PCR product bands obtained in different plants. (**b**) *AITR1* editing status in 21 individual T1 transgenic plants. PCR products were recovered from gel and sequenced. Sequencing results were examined and aligned with coding sequence of *AITR1* to check the editing status in the T1 transgenic plants. −, not edited, −/+, edited but heterozygous, ins, homozygous or biallelic editing with nucleotide insertions as indicated, N/A, not sequenced.

We therefore sequenced the full-length *AITR1* PCR products obtained from some of the plants that only produced one PCR product band, to see if we may get homozygous mutants with only one *AITR1* target site edited. Indeed, two of the early bolting plants, lines 11 and 15 were identified as homozygous mutants with a single nucleotide insertion at the second target site of *AITR1* (Figure 3b). One of the T1 plants with normal bolting phenotype was edited at the second site with a single nucleotide insertion, but was a biallelic mutant (Figure 3b). We also found that another 9 early bolting plants sequenced was edited in one of the target sites but were not homozygous (Figure 3b). It should note that among the single site edited mutant, only one was edited at the first target site (Figure 3b).

### Homozygous genome edited transgene-free mutants were obtained in the T2 plants

Having shown that homozygous mutants can be obtained in early bolting T1 transgenic plants (Figure 3), we decided to further examine whether transgene-free mutants can be easily obtained in T2 generation base on phenotypic observation. Two represent early mutant lines, i.e., line 14 and 15 were chosen, and T2 seeds were germinated directly in soil pots. Line 14 was chosen because PCR results indicated that both target sites of *AITR1* were edited in this line, where as line 15 was a homozygous mutant identified in T1 generation.

Segregation on bolting phenotype was observed in T2 plants (Figure 4a). Multiple plants without early bolting phenotype were obtained for both lines, and the segregation ratio for early and normal bolting plants were about 3:1 (Figure 4b).

**Figure 4.**
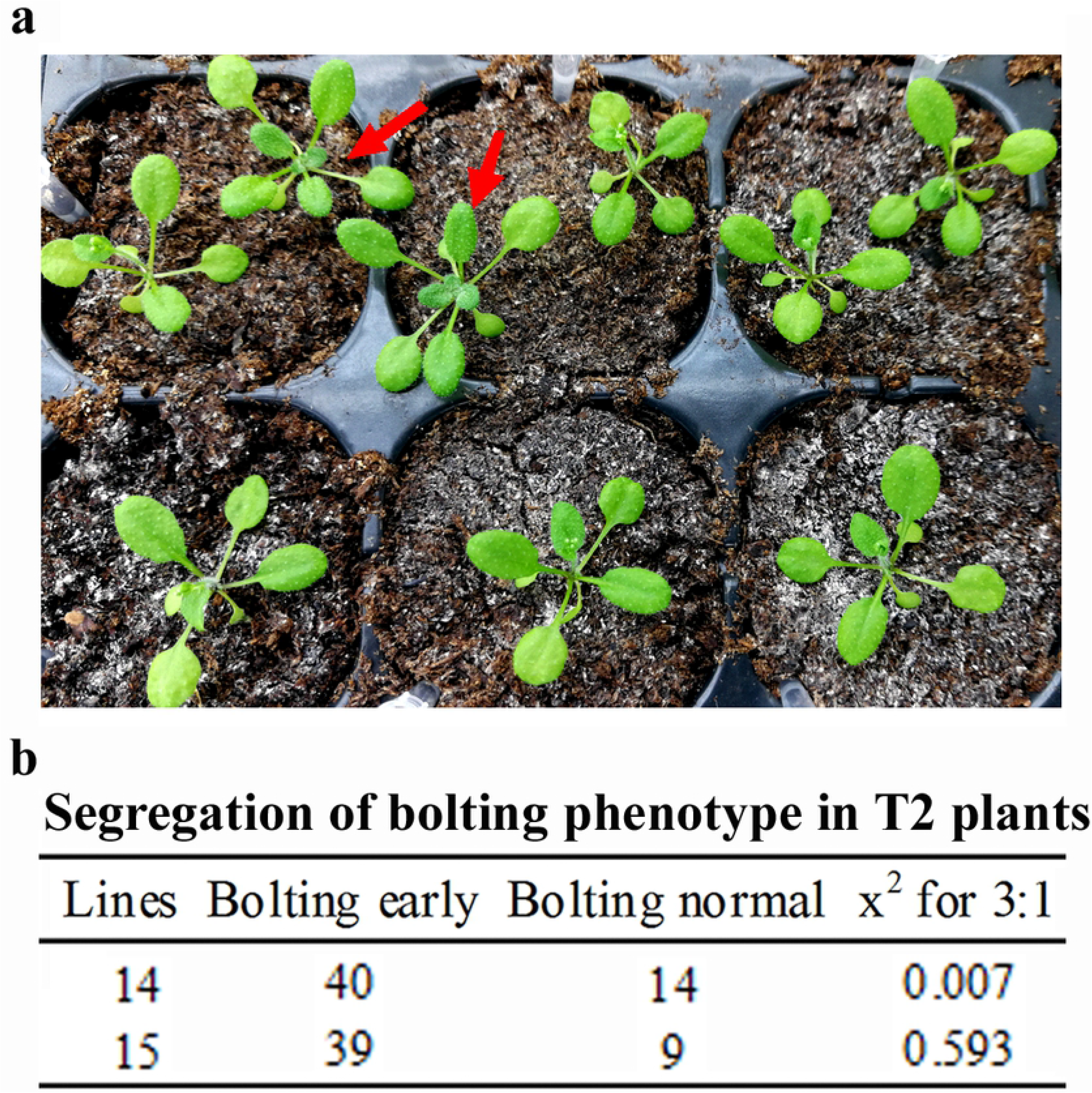
Phenotype segregation in the T2 generation of selected transgenic lines. (**a**) Bolting phenotype of the T2 plants from a single T1 transgenic line. T2 seeds were collected from selected T1 transgenic lines and sown directly into soil pots and grown in a growth room. Col wild type plants were generated and grown side by side with the T2 plants as a control. Pictures were taken 17 days after germination. Arrows indicate plants that were not bolting early. (**b**) Bolting phenotype segregation of the T2 plants from selected T1 transgenic lines. Chi square analysis was performed on omni calculator (https://www.omnicalculator.com/statistics/chi-square).

We then examined if the plants without early bolting phenotype were transgene-free plants by amplification of *Cas9* gene. As shown in Figure 5, no PCR products were obtained in all the normal bolting plants examined, whereas PCR products were obtained in early bolting plants. These results suggest that transgene-free plants can be easily identified by observation of bolting phenotype.

**Figure 5.**
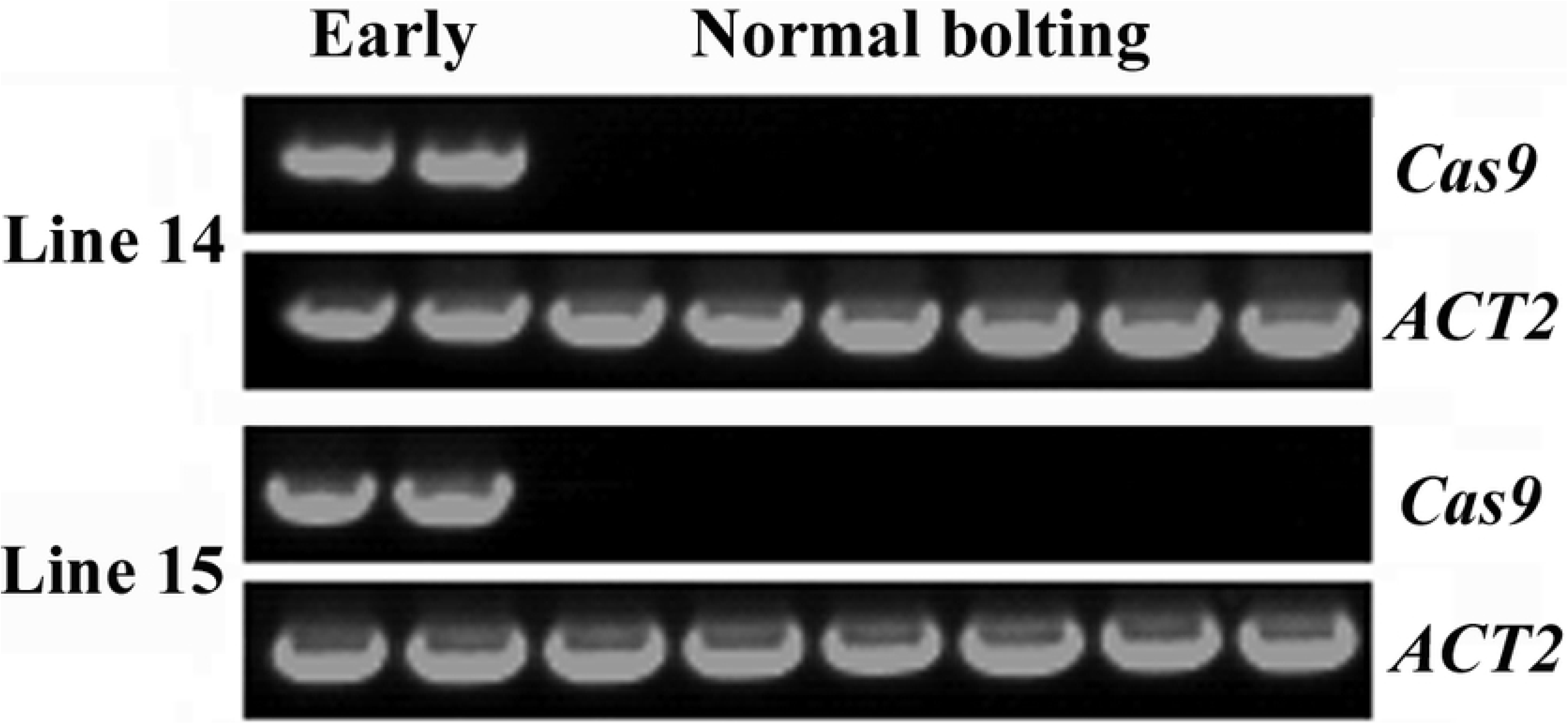
T2 plants with normal bolting time are transgene-free plants. DNA was isolated from leaves collected from six individual T2 plants with normal bolting time, and PCR was used to amplify *Cas9* fragment. For each line, DNA was also isolated from two early bolting plants, and used a positive control for *Cas9* amplification. PCR amplification of *ACT2* was used as a control.

By using PCR amplification, we found that two of the normal flowering plants in line 14 produced only a small band (Figure 6a), indicating that they were homozygous mutants. Sequence result showed that a 428bp fragment in *AITR1* was deleted in this mutant, leading to a few amino acid substitutions and premature stop after the 89^th^ amino acid residue (Figure 6a).

**Figure 6.**
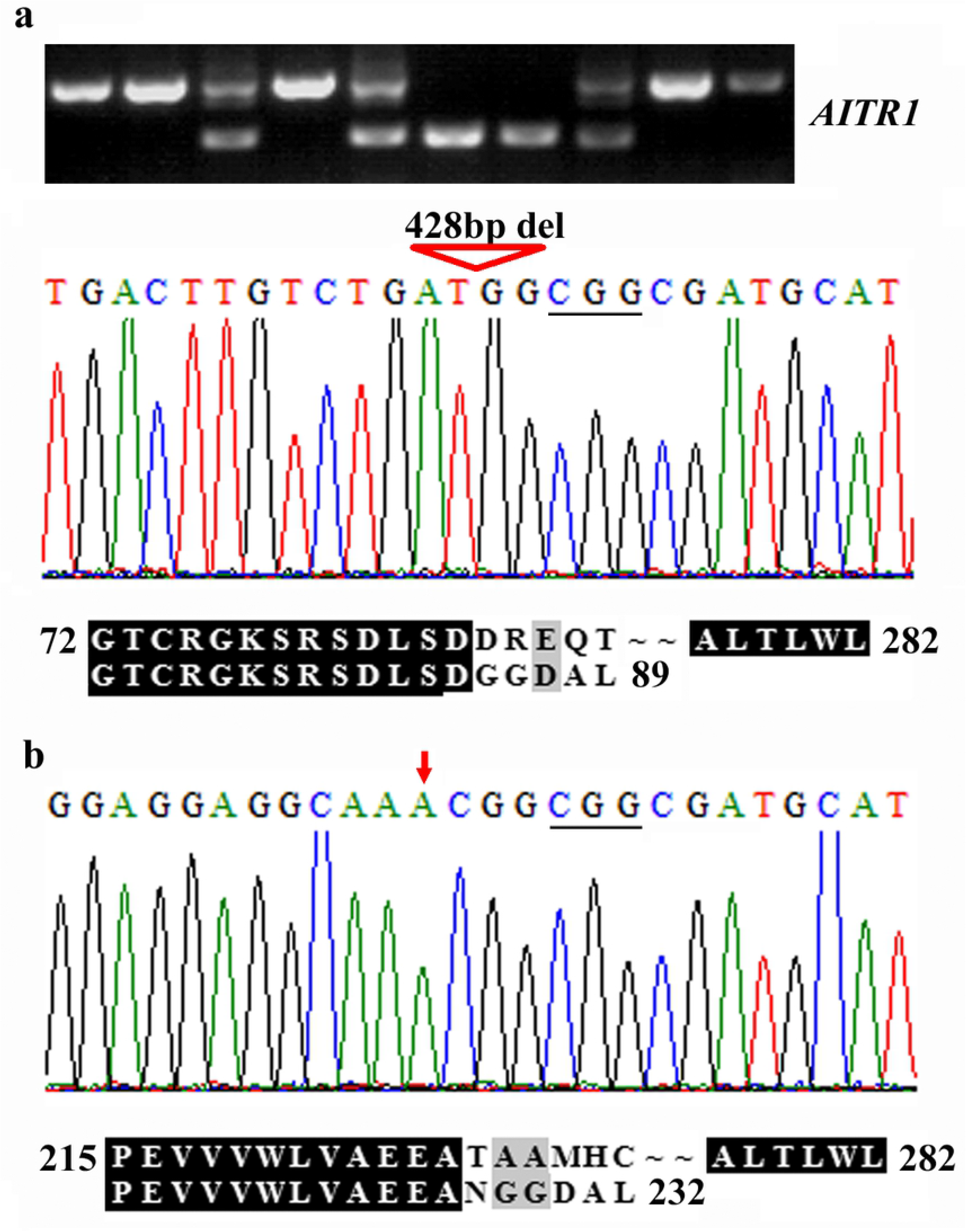
Isolation of genome edited transgene-free *aitr1* mutants. Transgene-free *aitr1* mutants isolated from line 14 (**a**) and line 15 (**b**). DNA was isolated from leaves collected from individual T2 plants with normal bolting time, and used as template to amplify the coding sequence of *AITR1*. The PCR products were recovered from gel and sequenced. Sequencing results were compared with coding sequence of *AITR1* to check the editing status. ORF of the *AITR1* sequences in the *aitr1* mutants were identified on ORFfinder (https://www.ncbi.nlm.nih.gov/orffinder/), and corresponding amino acid sequences were used for alignment with AITR1 amino acid sequences. Triangle indicates fragment deletion, arrow indicates nucleotide insertion, underlines in the sequence results indicate the PAM sites. Numbers in the alignment indicate the position of amino acid relative to the first Met of AITR1.

We also sequenced AITR PCR product from one of the transgene-free plants obtained from line 15 to confirm the genome editing status. We found that it was indeed a homozygous mutant with a single nucleotide insertion occurred in the sequence of the second target site. In this mutant, the single nucleotide insertion *AITR1* also led to a few amino acid substitutions, and premature stop occurred after the 232^nd^ amino acid residue (Figure 6b).

## Discussion

CRISPR/Cas9 genome-editing has been used to generate mutants in different plant species [6–12], and may have a bright future in using for plant breeding [6,12–14]. However, remove of the Cas9 T-DNA from the transgenic plants is likely necessary to get stable and heritable mutants, expecially for commercial use of the genome-edited crops [7,10].

FT promotes flowering in many plant species including the model plants Arabidopsis, crops such as rice and soybean, and fruit trees such as apple and pear [24–27,33], and has been used to accelerate plant breeding process by reducing juvenile phase of many plants [15,27,29].

By inserting the *GmFT2a* expression cassette into the *pHEE* CRISPR/Cas9 vector (Figure 1), we established a method for fast generation and easy identification of genome-edited transgene-free mutants in Arabidopsis (Figure 7).

**Figure 7.**
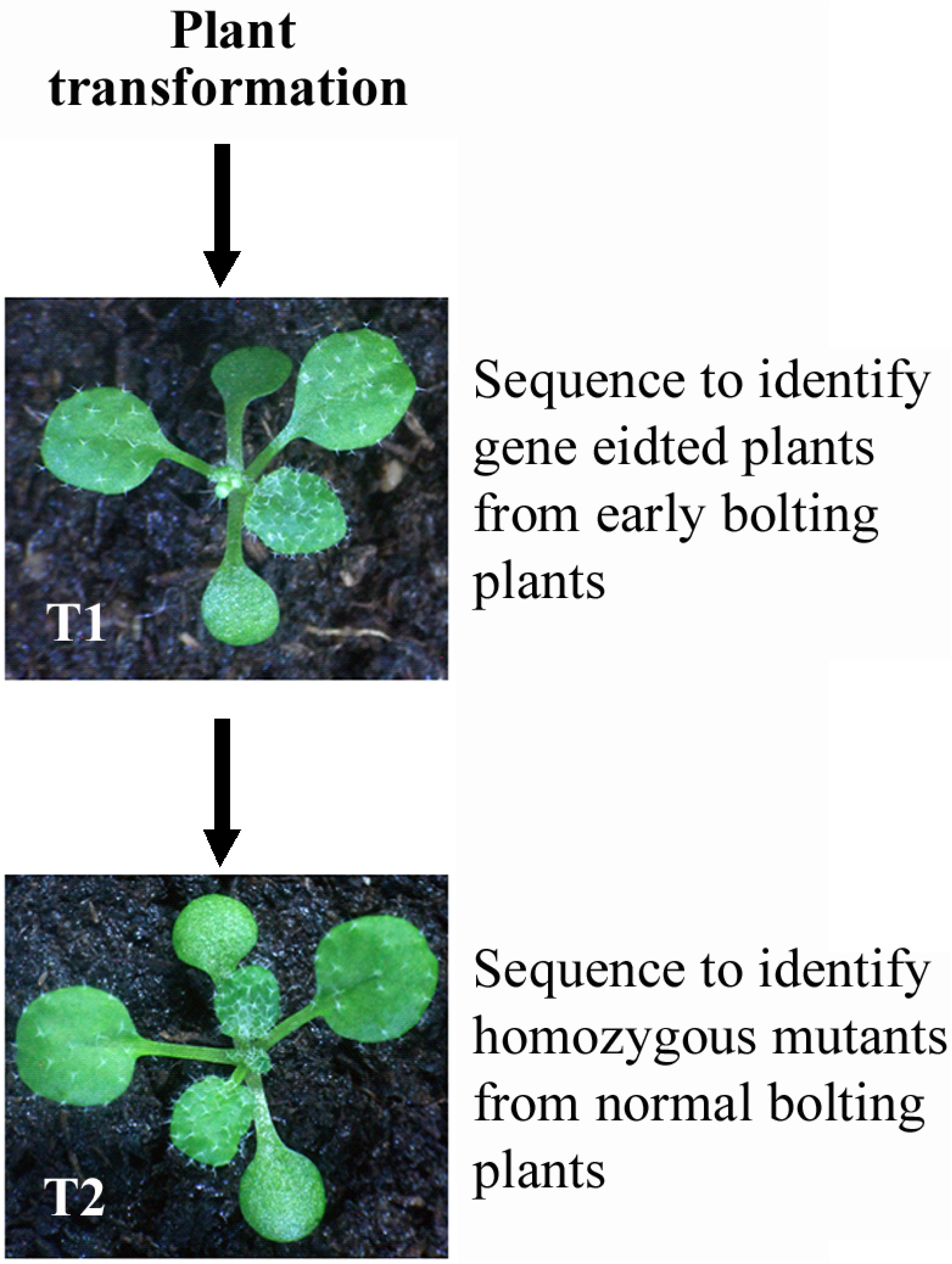
Simplified procedure for generating genome edited tansgene-free mutants by using *FT* expression cassette-containing CRISPR/Cas9 construct. Plants can be transformed and transgenic plants can selected in a way similar to that for other constructs. Within the T1 transgenic plants, select plants with early-bolting phenotype, and sequence to examine genome editing status. Keep only genome edited plants. In the T2 plants germinated from seeds of the individual T1 plants, keep only these bolting normally, and sequence to identify genome edited plants. These plants are genome edited transgene-free mutants. If necessary, confirm the transgene-free status by PCR amplification of *Cas9* fragment.^r^¡gure 1Bolting time (Days after transfer)

In this method, the easy visible early flowering phenotype can be used as an indicator of plants with Cas9 T-DNA. In the T1 generation, only plants with early bolting phenotypes should be selected and genome editing status should be examined to identify mutations. In the T2 generation of the genome-edited T1 plants, only plants without early flowering phenotypes, i.e., transgene-free plants should be selected, and genome editing status should be examined to ensure that genome-edited transgene-free homozygous mutants will be obtained (Figure 7).

By using the *pHEE-FT* vector to generate a CRISPR/Cas9 genome editing construct for simultaneously targeting two sites in the ABA signaling and abiotic stress tolerance regulator gene *AITR1* [30], we successfully obtained genome-edited transgene-free *aitr* homozygous mutants (Figure 6). In the T1 generation, about two thirds of the early bolting plants examined have at least one target site edited (Figure 3), indicating that insertion of the *GmFT2a* expression cassette into the CRISPR/Cas9 vector did not affect the editing efficiency of Cas9. However, we noted that including the two plants with fragment deletion, only three plants had mutations at the first target site, suggesting that the two targets selected have different editing efficiency. Had both target sites have high editing efficiency, we should able to obtained homozygous fragments deletion mutants in T1 generation. Nevertheless, we obtained homozygous mutants with mutations occurred at only one target site from the T1 plants (Figure 3), and we obtained homozygous fragments deletion mutants in T2 generations (Figure 6).

Even though only the double-stranded DNA cleave based CRISPR/Cas9 genome-editing system was examined in this study, however, the concept used in this study may also applied to the nucleotide substitution based CRISPR/Cas9 genome-editing system to facility transgene-free mutant isolation.

It should note that target site editing were also observed in T1 transgenic plants with medium or normal bolting time (Figure 3), suggesting that editing efficiency may not positively correlated with early bolting phenotypes. This is likely because *FT* and *Cas9* in the vector were in two different expression cassettes, and were driven by different promoters (Figure 1), thus their expression levels may not always positively related in the transgenic plants. Consider that mutants can obtained from early bolting T1 plants based on PCR results only or combined with sequencing (Figure 2), and transgene-free plants can be easily obtained from offsprings of the early bolting T1 plants based solely on bolting phenotype segregation (Figure 5), only T1 plants with early bolting phenotypes should be selected for next step analysis when using this method to generate for genome edited transgene-free mutants.

In addition to enable easy identification of transgene-free mutants base on phenotypic observation, the early bolting phenotype also reduce the length of the juvenile phase, which led to reduced overall time length required for generating genome edited transgene-free mutants. In our case, the different bolting time between early bolting transgenic plants and Col wild type plants were more than 10 days (Figure 2). Because early bolting plants were selected in T1, but plants with normal bolting time were selected in T2, the overall time length required for generating genome edited transgene-free mutants in Arabidopsis will be reduced at least 10 days. In some cases, transgene-free mutants may not be able to be identified in T2 generations, and had to be identified on T3 generations. In that case, identify mutant from offsprings of the gene-edited early bolting T2 plants may save more time than from that of the transgene-free heterozygous T2 mutants.

In this study, the effects of integration of *FT* expression cassette into CRISPR/Cas9 vector to accelerate transgene-free mutant isolation was tasted only in Arabidopsis, however, considering that the juvenile phase for most of the crops such as rice and soybean lasts for moths, and that for most of the trees including fruit trees apple and pear lasts for years, whereas overexpression of *FT* greatly reduced the length of juvenile phase in most of the plants examined, including all the plants mentioned above [25–28,34–36], the method described here may benefit even more for genome editing based breeding for the plants with a long juvenile phase.

On the other hand, transgenic plants of different plant species may need to be selected in different antibiotics, and multiple genes may need to be edited simultaneously, by replacing the antibiotic gene and/or the sgRNA clone cassette, the pHEE-FT vector reported here may be used for genome editing for a single or multiple genes in different plant species.

## Materials and methods

### Plant materials and growth conditions

The Columbia ecotype ‘Col-0’ (Col) Arabidopsis (*Arabidopsis thaliana*) was used for plant transformation, and as controls for bolting time assays.

Seeds of Col wild type were germinated in soil pots, and grown in a growth room. T1 transgenic plants were selected by plating T1 seeds on antibiotic-containing 1/2 MS plates. Transgenic seedlings were transferred into soil pots, and grown in a growth room. As a control, seeds of Col wild type were germinated on 1/2MS plates, seedlings were transferred into soil pots and grown in a growth room. The growth conditions in the growth room have been described previously [37,38].

### Constructs

The *pHEE* CRISPR/Cas9 vector has been described previously [33]. To insert the FT expression cassette into *pHEE* to generate the *pHEE-FT* vector, the full-length open reading frame (ORF) sequence of *GmFT2a* was synthesized and cloned into *pUC19* under the control of the double *35S* promoter, and terminated by *nos* [39]. The sequence of the *35S:GmFT2a-nos* cassette was then amplified by PCR, and cloned into the *pHEE* vector at the *Pme1* site by using Gibson assembly. The primers used to amplify *35S:GmFT2a-nos* from the *pUC19* construct were: 5’-CCTGTCAAACACTGATAGTTTGTCGACTCTAGAGGATCC-3’ and 5’-GTCGTTTCCCGCCTTCAGTTTACGACGGCCAGTGAATTC-3’.

To generate CRISPR/Cas9 construct for genome editing of *AITR1*, appropriate target sequences were first identified by scanning coding sequence of *AITR1* on CRISPRscan (http://www.crisprscan.org/?page=sequence), and then evaluated on Cas-OFFinder (http://www.rgenome.net/cas-offinder/). Two target sequences were selected and used for editing *of AITR1*, 5’-GATCTGACTTGTCTGATGAT(CGG)-3’, and 5’-GGTGGCGGAGGAGGCAACGG(CGG)-3’. The sgRNA expression cassettes targeting *AITR1* were cloned into the *pHEE-FT* vector to generate *pHEE-FT-AITR1* construct by following the procedures described by Wang et al [39]. The primers used to insert the target sequences into sgRNA expression cassettes were, *DT1-BsF*, 5’-ATATATGGTCTCGATTGATCTGACTTGTCTGATGATGTT-3’, *DTI-F0*, 5’-T GATCTGACTTGTCTGATGATGTTTTAGAGCTAGAAATAGC-3’, *DT2-R0*, 5-AACCCGTTGCCTCCTCCGCCACCAATCTCTTAGTCGACTCTAC-3’, and *DT2-BsR*, 5’-ATTATTGGTCTCGAAACCCGTTGCCTCCTCCGCCACC-3’. The primers used for colony PCR and to sequence the sgRNA expression cassettes in the generated *pHEE-FT-AITR1* construct were, *U6-26-IDF*, 5’-TGTCCCAGGATTAGAATGATTAGGC-3’ and *U6-26-IDR*, 5’-AGCCCTCTTCTTTCGATCCATCAAC-3’.

### Plant transformation and transgenic plant selection

Col wild type plants have several mature flowers on the main inflorescence (~5-week-old) were used for transformation. The plants were transformed via Agrobacterium *GV3101* mediated floral dipping [40]. T1 seeds collected were germinated on 1/2 MS plates containing 30 μg/ml hygromycin and 100 μg/ml carbenicillin to select transgenic plants.

Bolting time of the T1 plants was observed, and gene editing status was examined by amplifying the coding sequence of *AITR1* and sequencing the PCR products obtained.

### Isolation of transgene-free homozygous mutants

T2 seeds were collected from selected plants that bolting early and with *AITR1* gene edited. The seeds were then sown directly into soil pots. T2 plants with normal bolting time were selected, the absence of *Cas9* T-DNA insertion was confirmed by PCR amplification of *Cas9* fragment, and gene editing status was further examined/confirmed amplifying the coding sequence of *AITR1* and sequencing the PCR products obtained.

### DNA isolation and PCR

To examine gene editing status of *AITR1*, DNA was isolated from leaves of the transgenic plants, and the coding sequence of *AITR1* was amplified by PCR. PCR products was recovered from gel and sequenced. The sequencing results obtained were then examined and aligned with coding sequence of *AITR1* obtained from Phytozome (http://phytozome.jgi.doe.gov/pz/portal.html). The primers used for PCR amplification of *ACT2* and *AITR1* coding sequence and have been described previously [30,41].

To confirm the transgene-free status of the mutants obtained, DNA was isolated from leaves of T2 transgenic plants that have a normal bolting time, and *Cas9* gene fragment was amplified by PCR. The primers used for PCR amplification of *Cas9* were, *Cas9-F*, 5’-GGACAACGAGGAGAATGAGG-3’, and *Cas9-R*, 5’-TGTCTCGACCAGCTGCCTCTT-3’

### Bolting time assays

For the T1 plants, the date of bolting was recorded, and days after the transgenic seedlings were transferred into soil pots were calculated and used as bolting time. Transferred Col wild type plants were used as controls.

For T2 plants, the date of bolting was recorded, and days to bolting after the seeds were germinated were calculated. Col wild type plants germinated and grown in soil pots were used as controls.

## Author contribution statement

S.W. conceived the study. S.W., Y.C. and N.Z. designed the experiments. Y.C., N.Z., S.H., S.A., and W.Y. performed the experiments. S.W., Y.C. and N.Z. analyzed the data. S.W. drafted the manuscript. All the authors participated in the revision of the manuscript.

## Acknowledgement

This work was supported by the National Key R&D Program of China (2016YFD0101902). The funder has no role in study design, data collection and analysis, decision to publish, or preparation of the manuscript.

## References

1. Jinek, M.; Chylinski, K.; Fonfara, I.; Hauer, M.; Doudna, J.A.; Charpentier, E. A programmable dual-RNA-guided DNA endonuclease in adaptive bacterial immunity. Science 2012, 337, 816–821.

2. Cong, L.; Ran, F.A.; Cox, D.; Lin, S.; Barretto, R.; Habib, N.; Hsu, P.D.; Wu, X.; Jiang, W.; Marraffini, L.A.; Zhang, F. Multiplex genome engineering using CRISPR/Cas systems. Science 2013, 339, 819–823.

3. Li, J.F.; Norville, J.E.; Aach, J.; McCormack, M.; Zhang, D.; Bush, J.; Church, G.M.; Sheen, J. Multiplex and homologous recombination-mediated genome editing in Arabidopsis and Nicotiana benthamiana using guide RNA and Cas9. Nat Biotechnol. 2013, 31, 688–691.

4. Nekrasov, V.; Staskawicz, B.; Weigel, D.; Jones, J.D.; Kamoun, S. Targeted mutagenesis in the model plant Nicotiana benthamiana using Cas9 RNA-guided endonuclease. Nat Biotechnol. 2013, 31, 691–693.

5. Shan, Q.; Wang, Y.; Li, J.; Zhang, Y.; Chen, K.; Liang, Z.; Zhang, K.; Liu, J.; Xi, J.J.; Qiu, J.L., Gao, C. Targeted genome modification of crop plants using a CRISPR-Cas system. Nat Biotechnol. 2013, 31, 686–688.

6. Ma, X.; Zhang, Q.; Zhu, Q.; Liu, W.; Chen, Y.;, Qiu, R.; Wang, B.; Yang, Z.; Li, H.; Lin, Y.; Xie, Y.; Shen, R.; Chen, S.; Wang, Z.; Chen, Y.; Guo, J.; Chen, L.; Zhao, X.; Dong, Z.; Liu, Y.G. A robust CRISPR/Cas9 system for convenient, high-efficiency multiplex genome editing in monocot and dicot plants. Mol Plant 2015, 8, 1274–1284.

7. Gao, X.; Chen, J.; Dai, X.; Zhang, D.; Zhao, Y. An effective strategy for reliably isolating heritable and Cas9-free *Arabidopsis* mutants generated by RISPR/Cas9-mediated genome editing. Plant Physiol. 2016, 171, 1794–1800.

8. Lu, H.P.; Liu, S.M.; Xu, S.L.; Chen, W.Y.; Zhou, X.; Tan, Y.Y.; Huang, J.Z.; Shu, Q.Y. CRISPR-S: an active interference element for a rapid and inexpensive selection of genome-edited, transgene-free rice plants. Plant Biotechnol J. 2017, 15, 1371–1373.

9. Shimatani, Z.; Kashojiya, S.; Takayama, M.; Terada, R.; Arazoe, T.; Ishii, H.; Teramura, H.; Yamamoto, T.; Komatsu, H.; Miura, K.; Ezura, H.; Nishida, K.; Ariizumi, T.; Kondo, A. Targeted base editing in rice and tomato using a CRISPR-Cas9 cytidine deaminase fusion. Nat Biotechnol. 2017, 35, 441–443.

10. He, Y.; Zhu, M.; Wang, L.; Wu, J.; Wang, Q.; Wang, R.; Zhao, Y. Programmed Self-Elimination of the CRISPR/Cas9 Construct Greatly Accelerates the Isolation of Edited and Transgene-Free Rice Plants. Mol Plant 2018, 11, 1210–1213.

11. Zsögön, A.; Čermák, T.; Naves, E.R.; Notini, M.M.; Edel, K.H.; Weinl, S.; Freschi, L.; Voytas, D.F.; Kudla, J.; Peres, L.E.P. De novo domestication of wild tomato using genome editing. Nat Biotechnol. 2018, 36, 1211–1216.

12. Chen, K.; Wang, Y.; Zhang, R.; Zhang, H.; Gao, C. CRISPR/Cas Genome Editing and Precision Plant Breeding in Agriculture. Annu Rev Plant Biol. 2019, doi: 10.1146/annurev-arplant-050718-100049.

13. Hu, J.H.; Miller, S.M.; Geurts, M.H.; Tang, W.; Chen, L.; Sun, N.; Zeina, C.M.; Gao, X.; Rees, H.A.; Lin, Z.; Liu, D.R. Evolved Cas9 variants with broad PAM compatibility and high DNA specificity. Nature 2018, 556, 57–63.

14. Nishimasu, H.; Shi, X.; Ishiguro, S.; Gao, L.; Hirano, S.; Okazaki, S.; Noda, T.; Abudayyeh, O.O.; Gootenberg, J.S.; Mori, H.; Oura, S.; Holmes, B.; Tanaka, M.; Seki, M.; Hirano, H.; Aburatani, H.; Ishitani, R.; Ikawa, M.; Yachie, N.; Zhang, F.; Nureki, O. Engineered CRISPR-Cas9 nuclease with expanded targeting space. Science 2018, 361, 1259–1262.

15. Jung, C.; Muller, A.E. Flowering time control and applications in plant breeding. Trends Plant Sci. 2009, 14, 563–573.

16. Boss, P.K.; Bastow, R.M.; Mylne, J.S.; Dean, C. Multiple pathways in the decision to flower: enabling, promoting, and resetting. Plant Cell 2004, 16, S18-S31.

17. Yant, L.; athieu, J.; Schmid, M. Just say no: floral repressors help Arabidopsis bide the time. Curr Opin Plant Biol. 2009, 12, 580–586.

18. Wahl, V.; Ponnu, J.; Schlereth, A.; Arrivault, S.; Langenecker, T.; Franke, A.; Feil, R.; Lunn, J.E.; Stitt, M.; Schmid, M. Regulation of flowering by trehalose-6-phosphate signaling in Arabidopsis thaliana. Science 2013, 339, 704–707.

19. Gendall, A.R.; Levy, Y.Y.; Wilson, A.; Dean, C. The VERNALIZATION 2 gene mediates the epigenetic regulation of vernalization in Arabidopsis. Cell 2001, 107, 525–535

20. Ratcliffe, O.J.; Kumimoto, R.W.; Wong, B.J.; Riechmann, J.L. Analysis of the Arabidopsis MADS AFFECTING FLOWERING gene family: MAF2 prevents vernalization by short periods of cold. Plant Cell 2003, 15, 1159–1169.

21. Mockler, T.C.; Yu, X.H.; Shalitin, D.; Parikh, D.; Michael, T.P.; Liou, J.; Huang, J.; Smith, Z.; Alonso, J.M.; Ecker, J.R.; Chory, J.; Lin, C. Regulation of flowering time in Arabidopsis by K homology domain proteins. Proc Natl Acad Sci USA 2004, 101. 12759–12764.

22. Tamada, Y.; Yun, J.Y.; Woo, S.C.; Amasino, R.M. ARABIDOPSIS TRITHORAX-RELATED7 is required for methylation of lysine 4 of histone H3 and for transcriptional activation of FLOWERING LOCUS C. Plant Cell 2009, 21, 3257–3269.

23. Wang, J.W.; Czech, B.; Weigel, D. miR156-regulated SPL transcription factors define an endogenous flowering pathway in Arabidopsis thaliana. Cell 2009, 138, 738–749.

24. Jung, J.H.; Seo, P.J.; Ahn, J.H.; Park, C.M. Arabidopsis RNA-binding protein FCA regulates microRNA172 processing in thermosensory flowering. J Biol Chem. 2012, 287, 16007–16016

25. Tamaki, S.; Matsuo, S.; Wong, H.L.; Yokoi, S.; Shimamoto, K. Hd3a protein is a mobile flowering signal in rice. Science 2007, 316, 1033–1036.

26. Kong, F.; Liu, B.; Xia, Z.; Sato, S.; Kim, B.M.; Watanabe, S.; Yamada, T.; Tabata, S.; Kanazawa, A.; Harada, K.; Abe, J. Two coordinately regulated homologs of FLOWERING LOCUS T are involved in the control of photoperiodic flowering in soybean. Plant Physiol. 2010, 154, 1220–1231.

27. Putterill, J.; Varkonyi-Gasic, E. FT and florigen long-distance flowering control in plants. Curr Opin Plant Biol 2016, 33, 77–82.

28. Wickland, D.P.; Hanzawa, Y. The FLOWERING LOCUS T/TERMINAL FLOWER 1 gene family: functional evolution and molecular mechanisms. Mol Plant 2015, 8, 983–997.

29. Bull, S.E.; Seung, D.; Chanez, C.; Mehta, D.; Kuon, J.E.; Truernit, E.; Hochmuth, A.; Zurkirchen, I.; Zeeman, S.C.; Gruissem, W.; Vanderschuren, H. Accelerated ex situ breeding of GBSS- and PTST1-edited cassava for modified starch. Sci Adv. 2018, 4, eaat6086.

30. Tian, H.; Chen, S.; Yang, W.; Wang, T.; Zheng, K.; Wang, Y.; Cheng, Y.; Zhang, N.; Liu, S.; Li, D.; Liu, B.; Wang, S. A novel family of transcription factors conserved in angiosperms is required for ABA signalling. Plant Cell Environ. 2017, 40, 2958–2971.

31. Sun, H.; Jia, Z.; Cao, D.; Jiang, B.; Wu, C.; Hou, W.; Liu, Y.; Fei, Z.; Zhao, D.; Han, T. GmFT2a, a soybean homolog of FLOWERING LOCUS T, is involved in flowering transition and maintenance. PLoS One 2011, 6, e29238.

32. Nan, H.; Cao, D.; Zhang, D.; Li, Y.; Lu, S.; Tang, L.; Yuan, X.; Liu, B.; Kong, F. GmFT2a and GmFT5a redundantly and differentially regulate flowering through interaction with and upregulation of the bZIP transcription factor GmFDL19 in soybean. PLoS One 2014, 9, e97669.

33. Wang, Z.; Xing, H.; Dong, L.; Zhang, H.; Han, C.; Wang, X.; Chen, Q. Egg cell-specific promoter-controlled CRISPR/Cas9 efficiently generates homozygous mutants for multiple target genes in Arabidopsis in a single generation. Genome Biol. 2015, 16, 144.

34. Putterill, J.; Zhang, L.; Yeoh, C.; Balcerowicz, M.; Jaudal, M.; Varkonyi Gasic, E. FT genes and regulation of flowering in the legume Medicago truncatula. Funct Plant Biol. 2013. 40, 1199–1207

35. Yamagishi, N.; Kishigami, R.; Yoshikawa, N. Reduced generation time of apple seedlings to within a year by means of a plant virus vector: a new plant-breeding technique with no transmission of genetic modification to the next generation. Plant Biotechnol J. 2014, 12. 60–68.

36. Yamagishi, N.; Li, C.; Yoshikawa, N. Promotion of flowering by Apple latent spherical virus vector and virus elimination at high temperature allow accelerated breeding of apple and pear. Front Plant Sci. 2016, 7, 171.

37. Tian, H.; Guo, H.; Dai, X.; Cheng, Y.; Zheng, K.; Wang, X.; Wang, S. An ABA down-regulated bHLH transcription repressor gene, bHLH129 regulates root elongation and ABA response when overexpressed in Arabidopsis. Scic Rep. 2015, 5, 17587.

38. Dai, X.; Zhou, L.; Zhang, W.; Cai, L.; Guo, H.; Tian, H.; Schiefelbein, J.; Wang, S. A single amino acid substitution in the R3 domain of GLABRA1 leads to inhibition of trichome formation in Arabidopsis without affecting its interaction with GLABRA3. Plant Cell Environ. 2016, 39, 897–907.

39. Wang, S.; Tiwari, S.B.; Hagen, G.; Guilfoyle, T.J. AUXIN RESPONSE FACTOR7 restores the expression of auxin-responsive genes in mutant Arabidopsis leaf mesophyll protoplasts. Plant Cell 2005, 17, 1979–1993.

40. Clough, S.J.; Bent, A.F. Floral dip: a simplified method for Agrobacterium-mediated transformation of Arabidopsis thaliana. Plant J. 1998, 16, 735–743.

41. Guo, H.; Zhang, W.; Tian, H.; Zheng, K.; Dai, X.; Liu, S.; Hu, Q.; Wang, X.; Liu, B.; Wang, S. An auxin responsive CLE gene regulates shoot apical meristem development in Arabidopsis. Front Plant Sci. 2015, 6, 295.

